# Partial recurrence enables robust and efficient computation

**DOI:** 10.1101/2025.07.28.667142

**Authors:** Marcus Ghosh, Dan F. M. Goodman

## Abstract

Neural circuits are sparse and bidirectional. Meaning that signals flow from early sensory areas to later regions and back. Yet, between connected areas there exist some but not all pathways. How does this structure, somewhere between feedforward and fully recurrent, shape circuit function? To address this question, we designed a recurrent neural network model in which a set of weight matrices (i.e. pathways) can be combined to generate every network structure between feedforward and fully recurrent. We term these architectures partially recurrent neural networks (pRNNs). We trained over 25,000 pRNNs on a novel set of reinforcement learning tasks, designed to mimic multisensory navigation, and compared their performance across multiple functional metrics. Our findings reveal three key insights. First, many architectures match or exceed the performance of fully recurrent networks, despite using as few as one-quarter the number of parameters; demonstrating that partial recurrence enables energy efficient, yet performant solutions. Second, each pathway’s functional impact is both task and circuit dependent. For instance, feedback connections enhance robustness to noise in some, but not all contexts. Third, different pRNN architectures learn solutions with distinct input sensitivities and memory dynamics, and these computational traits help to explain their functional capabilities. Overall, our results demonstrate that partial recurrence enables robust and efficient computation - a finding that helps to explain *why* neural circuits are sparse and bidirectional, and *how* these principles could inform the design of artificial systems.

**1 Key Points:**

- We define a neural network model in which a set of weight matrices (i.e. pathways) can be combined to generate every network structure between feedforward and fully recurrent.
- Many of these partially recurrent (pRNN) architectures perform as well as, or even better than, fully recurrent networks, in terms of their task performance, sample efficiency and robustness to various perturbations.
- Each pathway’s functional impact is both task and circuit dependent. For example, input to output skip connections boost learning speed in some, but not all tasks or architectures.
- We propose a conceptual model, whereby: networks with different structures learn solutions with distinct computational traits, which shape their function.

## 2 Introduction

Across species, anatomical studies reveal a striking feature of neural circuits: they are sparse, yet bidirectional. At the level of single neurons, the *C*.*elegans* and *Drosophila* connectomes contain only ≈3% (White et al., 1986; Cook et al., 2019; Witvliet et al., 2021) and *<*1% (Winding et al., 2023; Schlegel et al., 2024; Lin et al., 2024) of possible synapses, yet an abundance of feedback connections. Similarly, at the level of region-region pathways in the mouse (Oh et al., 2014; Zingg et al., 2014) and macaque neocortex (Markov et al., 2014) only ≈50% of areas connect, yet many of these connections are reciprocal.

This anatomical structure raises a fundamental question: *what advantages or disadvantages do sparse, bidirectional circuits confer*, compared to either: feedforward networks – in which information simply flows from early sensory areas to later areas, or fully recurrent networks – in which every component is bidirectionally connected? Determining this is challenging *in vivo*, but *in silico* one can design networks with distinct architectures and compare different measures of function. Though, how should we explore the vast space of possible network structures?

Computational studies have primarily focussed on either feedforward (Yamins et al., 2014) or recurrent architectures (Mante et al., 2013; Sussillo et al., 2015). Though, three lines of work have imbued recurrent networks with properties closer to those found in real circuits. One approach is to impose a low-rank structure on the recurrent (hidden-hidden) weight matrix, effectively reducing its dimensionality by introducing structured correlations between units. This low-rank structure can be imposed prior to training (Liu et al., 2024), or even analytically derived to implement specific task-relevant dynamics (Mastrogiuseppe and Ostojic, 2018). However, as networks with very distinct structures can share the same rank, and altering connection weights changes the rank, but not the topology, interpreting structure-function relations in low-rank networks remains challenging (section 6.1). Another approach, in task-trained models, is to add regularization terms to the loss function, to encourage networks to learn recurrent connections with specific properties, such as modularity or small-worldness (Achterberg et al., 2023). Though, as in the low-rank case (Liu et al., 2024), these structural changes do not lead to changes in network function, defined as task performance (Achterberg et al., 2023) (with the exception of Zhang et al. 2025). Alternatively, one can train networks with sparse or modular recurrent weight matrices. One approach is to directly impose this structure (Goulas et al., 2021; Béna and Goodman, 2025). Another is to use learning algorithms which add and remove connections, throughout training, to achieve a given sparsity (Bellec et al., 2017; Jayakumar et al., 2020). Sparse networks are more energy efficient than dense networks, but this often comes with at least a small cost to performance in task-trained models (Bellec et al., 2017; Jayakumar et al., 2020; Goulas et al., 2021; Béna and Goodman, 2025). However, in other contexts, sparsity has been shown to support robust associative memory (Hoffmann, 2019), suggesting benefits beyond efficiency.

Taken together, functional results from these three approaches suggest that either many structures are degenerate, in that they yield the same input-output mappings (Edelman and Gally, 2001; Marder and Taylor, 2011), or that unconstrained networks, with more parameters, would always be more performant. However, it is possible that more naturalistic tasks or alternative measures of function, for example sample efficiency (i.e. learning speed) or robustness to noise, would reveal differences between these structures, and perhaps even scenarios in which sparser networks outperform dense networks.

Here, we explore the link between network structure and function in four steps. First, we introduce a set of multisensory maze tasks. In these tasks, we simulate networks as agents with noisy sensors, which provide clues about the shortest path through each maze. We focus on these tasks, as they involve translating the inputs from multiple sensory channels, *±* internal states, to actions. As such, these tasks more closely resemble the kinds of challenges animals face, than many machine learning tasks. Moreover, there are well established theoretical models of multisensory integration (Jones, 2016; Ghosh et al., 2024) which we use to benchmark our tasks. Second, we define a recurrent neural network model in which signal propagation is bidirectional, yet sparse. We term these models partially recurrent neural networks (pRNNs). Third, we train all pRNN architectures on our maze tasks, and compare their performance, sample efficiency and robustness to various perturbations. Finally, to link each architecture’s structure to its function, we measure three computational traits from each network: their weights, input sensitivity and memory dynamics, and quantify how predictive these traits are of network function. Our results demonstrate that partial recurrence enables robust and efficient computation, and explain why networks with distinct structures function differently.

## 3 Results

### 3.1 General setup

We explore a range of scenarios in which agents must navigate multisensory mazes. All agents operate in discrete time, taking fixed-width steps on a grid. For implementation details please see section 5.3.

In these tasks, we provide sensory cues in the form of positive spatial gradients leading towards the goal. While these gradients may seem abstract, they represent some stimuli well – such as chemical, thermal or luminance gradients. And, they provide a good approximation to the information provided by more complex cues. For example, tracks through the snow or mud which would appear crisper as one neared their source.

To perform these tasks, each agent implements a three-step sensation-action loop (Figure 1A). First, they sense their local environment using sensors from two channels, such as vision and hearing. Often, we add independent Gaussian noise to each channel at each time step. Next, they implement a policy mapping their sensory inputs to potential actions. Finally, they choose and implement an action; either pausing or moving in a cardinal direction.

**Figure 1:**
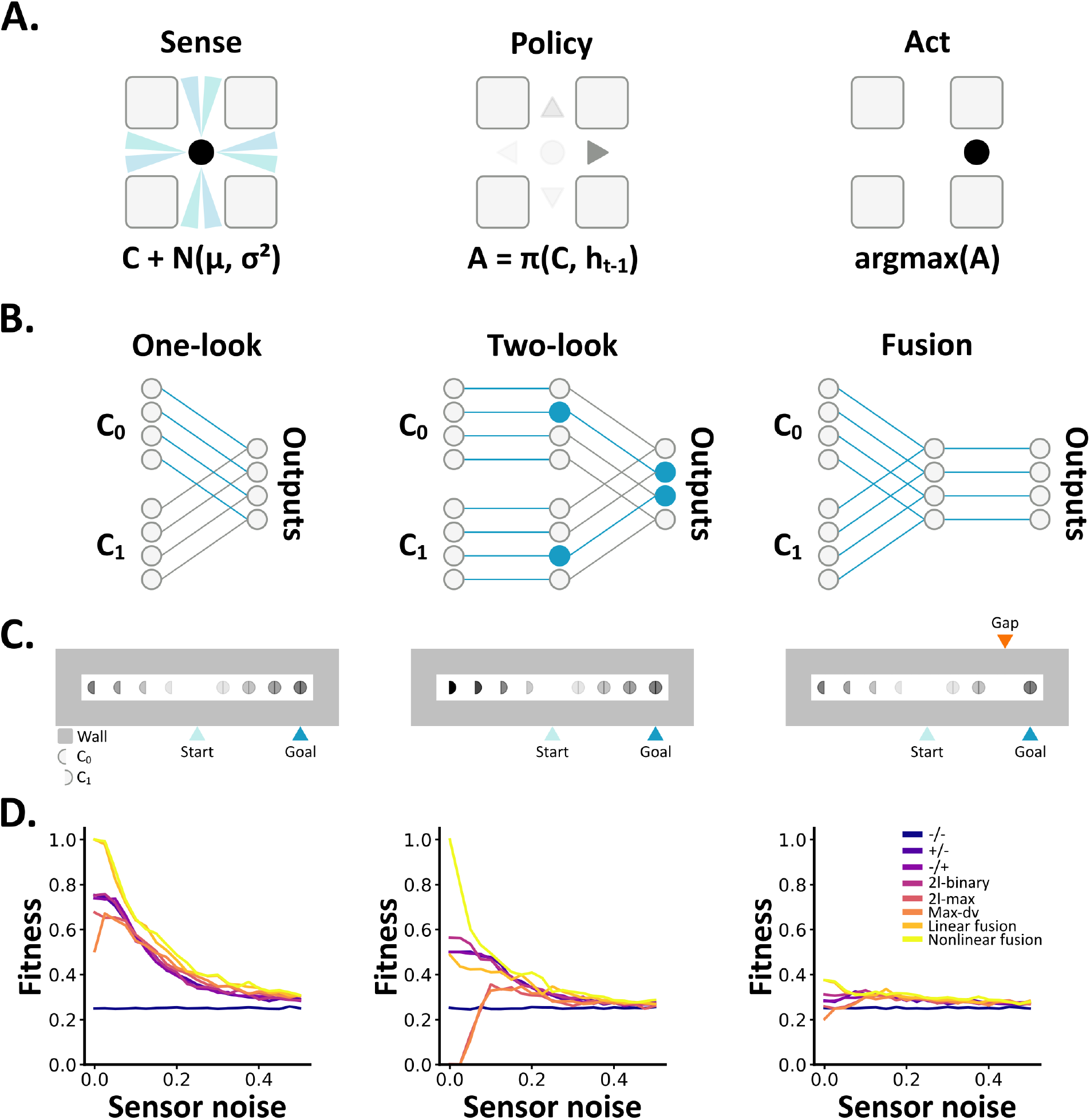
**A** In our tasks, agents must translate their noisy sensations (*C* + *N*(*µ, σ*^2^)), and in some cases hidden states (*h*_*t −* 1_), into actions (*A*) using algorithmic or learned policies (*π*). **B** Multisensory algorithms can solve these tasks by translating their sensory inputs (*C*) to actions (*A*) via different types of policies (section 5.4). One-look algorithms (left) use one channel (blue connections) but not the other (grey connections). Two-look algorithms (middle) make independent decisions per channel (blue circles) then base their output on these. Fusion algorithms form their outputs by combining their inputs across sensory channels. **C** Left: an example track maze with a path shown in white and walls in grey. Sensory cues are shown as circles, with cues from channel 0/1 shown on the left/right of each circle; such that semicircles denote unisensory cues. Light to dark fills indicate weak to strong signals. Middle: an example nonlinear track maze – the strength of the unisensory cues (leading away from the goal) matches the strength of the multisensory cues (leading towards). Right: an example track maze with a gap in the sensory cues. **D** Each algorithm’s mean fitness as a function of increasing sensor noise on the: track maze (left), nonlinear track maze (middle) and track maze with a gap (right).

### 3.2 Optimal multisensory algorithms are brittle

To benchmark our tasks, we began by testing how well seven classical models of multisensory integration (Figure 1B), interpreted as policies (section 5.4), perform a simple maze task.

In this task, (section 5.2.1), agents start in the middle of a 1D track. Their aim is to reach a goal location at either the left or right end of the track within a fixed number of time steps. To determine which end to navigate to, agents must use the signals from their sensors, which are grouped into two channels (C_0_ and C_1_). Multisensory signals, from both channels (C_0_ and C_1_), lead towards the goal. While, unisensory signals, from one channel (C_0_ or C_1_), lead away from the goal. Across trials, we vary both the goal location and which channel provides the unisensory signal. As the multisensory signal is twice as strong as the unisensory signal, the optimal strategy is to sum the information across channels and head in the direction with the greatest total.

As a measure of performance we define a fitness function from two metrics (section 5.6):

- Speed - the time taken to reach the goal. Normalised from 0 (not reached within a fixed number of time steps) to 1 (reached as quickly as possible).
- Accuracy - proximity to the goal after a fixed number of time steps. Normalised from 0 (ending the trial at the furthest point from the goal) to 1 (goal reached).

We weight speed and accuracy equally, such that fitness ranges from 0 to 1 (always taking the shortest path from start to goal). Notably, for consistency with later tasks, agents sense in 2D and can try to move up or down - in which case they will collide with the wall and remain in place.

In principle, we could learn parameters for these algorithms. However, as each sensor is equally reliable, we simply treat them as non-learning policies, which we do not train or fit in any way. Intuitively, agents which ignore both channels (-/-), and so move randomly, perform poorly and are unaffected by increasing sensor noise (Figure 1D, left). Agents which use one channel (+/- and -/+) or consider both but base their decision on only one (2l-binary, 2l-max, Max-dv) perform reasonably well and decay with noise (Figure 1D, left). Agents which fuse information across channels either linearly or nonlinearly perform optimally (Figure 1D, left), demonstrating the potential for tasks like these to reveal behavioural differences between models. To explore this further, we tested the same seven algorithms on two variants of this task.

First, we modified the cues by matching the strength of the unisensory signal leading away from the goal, to the multisensory signal leading towards it (Figure 1C, middle) (section 5.2.1). Consequently, at the first time step, a linear sum across channels is uninformative, as the total amount of signal on the agent’s left and right is equal (in the absence of noise). Instead, the optimal strategy is to use a nonlinear function to weight the multisensory signal more highly than the unisensory signal. As expected, and akin to Ghosh et al. (2024), nonlinear fusion now significantly outperforms the other algorithms (Figure 1D, middle). Strikingly, two algorithms (2l-max and Max-dv) even perform below chance (-/-), at low noise levels, as they base their decisions on only the strongest cue in either channel – which is misleading in this case.

Second, we introduced a gap in the sensory cues leading towards the goal (Figure 1C, right). Now, all seven algorithms perform near chance (-/-), as just prior to the gap they sense nothing ahead but something behind so, fearing the void, turn back (Figure 1D, right).

Together, these results highlight that these classic algorithms are brittle. In the sense that, while they are optimal in specific settings, relatively small changes can reduce them to chance performance or even mislead them entirely. This brittleness motivates our search for alternative, more robust multisensory models.

### 3.3 Partially recurrent neural networks

To explore alternative models, we began with a neural network with: a layer of input sensors (from two channels), a hidden layer and an output layer (with a node per action: move left, right, up, down or pause). Units in the input, hidden and output layers (respectively) use linear, rectified linear (ReLU) and softmax activation functions.

In this network, there are 9 possible connection pathways or weight matrices (Figure 2A):

**Figure 2:**
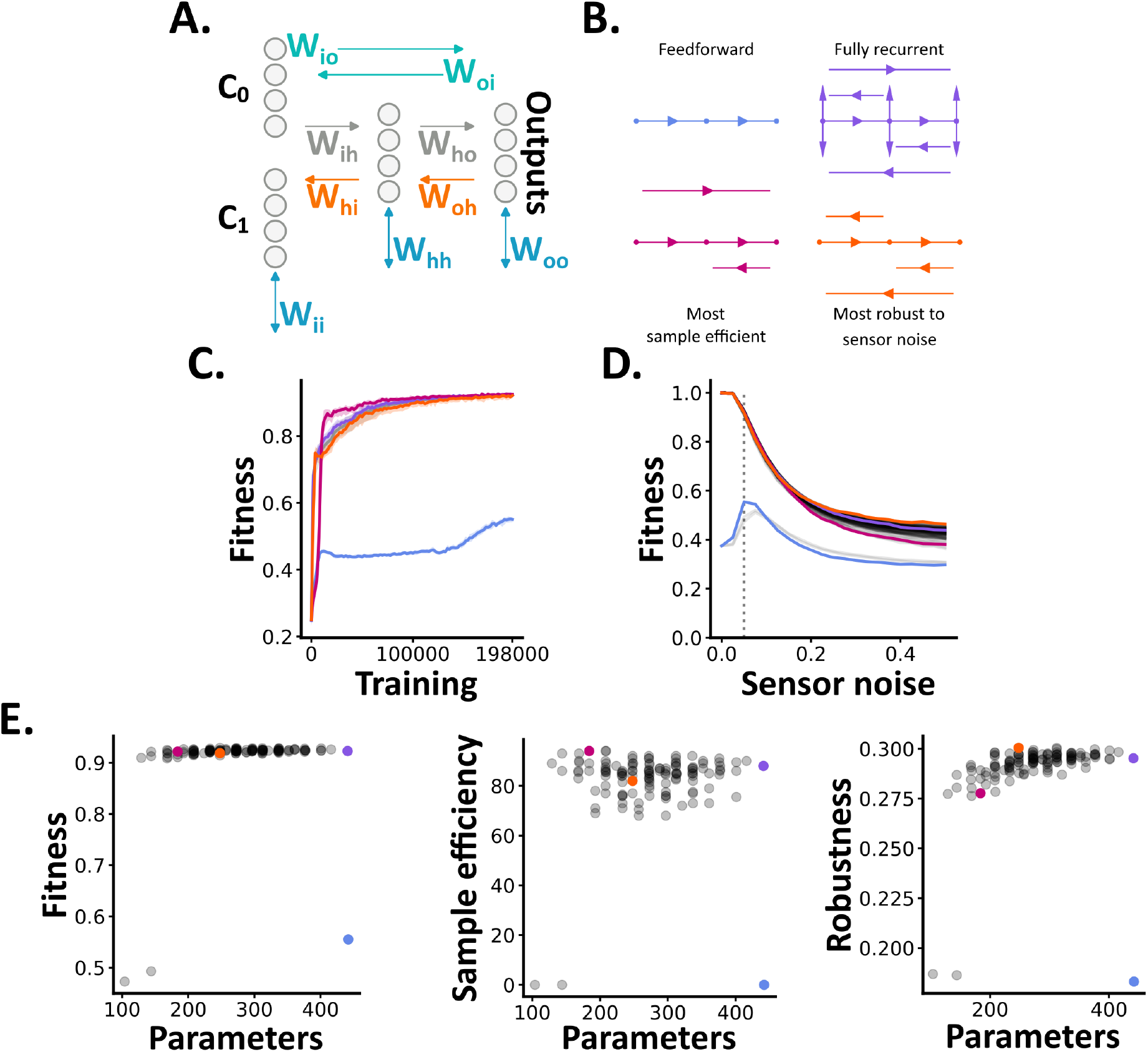
**A** Partially recurrent neural networks. Each architecture is composed of: input (i), hidden (h) and output units (o), two feedforward weight matrices *W*_*ih*_ and *W*_*ho*_ and any combination of the other 7 pathways. **B** 4 example architectures. **C** Test fitness across training on the track maze task, with a gap, for the 4 architectures shown in B (with matching colours), and all trained networks (grey). For each we plot the median (solid line) and interquartile range (shaded surround). Note that here, and for subsequent panels, feedforward denotes the large-feedforward architecture. **D** Each architecture’s median fitness as a function of increasing sensor noise on the track maze task with a gap. All architectures are drawn in grey, with those from panel B highlighted. The dashed vertical line denotes the level of sensor noise during training. **E** Each architecture’s number of trainable parameters vs median: fitness (left), sample efficiency (middle) and robustness to sensor noise (right). Each architecture is shown as a grey circle, with the architectures in B highlighted.

- **Feedforward** connections – from the input to hidden layer (*W*_*ih*_) and hidden to output layer (*W*_*ho*_).
- **Lateral** connections – connecting each unit to itself and the other units in its layer: input-input (*W*_*ii*_), hidden-hidden (*W*_*hh*_) and output-output (*W*_*oo*_).
- **Skip** connections – connecting the inputs directly to the outputs (*W*_*io*_) or vice versa (*W*_*oi*_).
- **Backwards** connections – connecting the hidden units to the inputs (*W*_*hi*_) or the output units to the hidden units (*W*_*oh*_).

Assuming the feedforward connections (*W*_*ih*_ and *W*_*ho*_) are always present, and each of the other 7 matrices can be present in any combination yields 2^7^ = 128 distinct architectures. In this framework, excluding all 7 additional weight matrices yields a pure layer-layer feedforward network (Figure 2B). While, including all 7 generates a fully recurrent network – as every unit connects bidirectionally to every other unit (Figure 2B). Between these two extremes are 126 architectures (e.g. Figure 2B).

We term these architectures **partially recurrent** as information flow can be *bidirectional* – from early to later areas and back, yet *sparse* – in the sense that it cannot flow via all possible paths, as in a fully recurrent network. While several of these architectures have been studied individually, for instance Jordan (1986) and Elman (1990) define networks with output to hidden connections (*W*_*oh*_) or lateral hidden connections (*W*_*hh*_) respectively. Our contribution is to place these, and other architectures, into a taxonomy, which provides a common reference frame for comparing different architectures and highlighting under-explored structures.

All 128 circuits are shown with their sparse layer-layer pathways or sparse unit-unit matrices in section 6.2. As we move through this space of network structures, how does function change? For instance, do networks with lateral hidden connections (*W*_*hh*_) – as in the standard recurrent neural network model, behave differently from those with lateral input (*W*_*ii*_) or output (*W*_*oo*_) connections? And, how would adding forward skip connections (*W*_*io*_) to those architectures change their behaviour?

### 3.4 Partial recurrence enables efficient, yet performant solutions

We began by training all 128 architectures on the track maze task with a gap (Figure 1C, right) using relatively small networks with 8 hidden units. To train each network’s weights we used reinforcement learning, specifically deep Q-learning (Mnih et al., 2013; Jensen, 2024). At each time step, pRNN agents act based on their current network output, without iterating to a steady state. To capture intra-architecture variability, across random seeds (Patterson et al., 2023), and to enable fair inter-architecture comparisons we trained 50 networks per architecture; yielding 6,400 trained networks.

Fully recurrent networks learned the task well, achieving a median fitness of 0.92 *±* 0.01 std (Figure 2C). By contrast, only two architectures were consistently unable to learn the task well: the pure layer-layer feedforward architecture (0.47 *±* 0.02 std), and an architecture with additional forward skip connections *W*_*io*_ (0.49 *±* 0.02 std). Notably, larger feedforward networks, with the same number of learnable parameters as the fully recurrent networks, were still unable to learn the task well (0.56 *±* 0.01 std)(Figure 2C). This is due to the fact that neither of these architectures retain any information from prior time steps, and so cannot solve tasks which require some form of memory. Strikingly, the remaining 125 architectures all achieved fitnesses close to the fully recurrent network (with median values between 0.91 and 0.93); despite having distinct pathways and far fewer learnable parameters - in some cases as low as ^1^*/*_4_ of the fully recurrent network (Figure 2E, left). While this may not be surprising, given the simplicity of the task, what is perhaps more surprising is that 16 architectures learned significantly faster than the fully recurrent network (Figure 2C and E middle). And, 9 were significantly more robust to noise (Figure 2D and E right). The fastest learning and most robust architectures are shown in fig. 2B.

These results demonstrate that many architectures can implement robust behaviours; equivalent, or even better than a fully-connected network. Though, would this hold true in more complex tasks?

To test this, we designed a 2 parameter family of multisensory mazes. Specifically, we create maze structures (walls and paths) using the Aldous-Broder algorithm (Aldous, 1990; Broder, 1989) and fill these with sensory gradients leading towards the goal (Figure 3A). This allows us to consider the more complex case where agents must navigate in four directions. Then, we introduce additional complexity by removing some fraction of walls – creating loops and open spaces (Figure 3A), and some fraction of sensory cues – creating gaps as in the track maze (Figure 3A). We focus on three scenarios: mazes with all walls and cues (M), mazes with 10% of walls removed (M_sw_) and mazes with 10% of cues removed (M_sc_). For each of these scenarios, we trained all 128 architectures, plus a large feedforward network, with 50 repeats per architecture. Combined with the networks from the track task with a gap (T_sc_) this yields over 25,000 trained networks.

**Figure 3:**
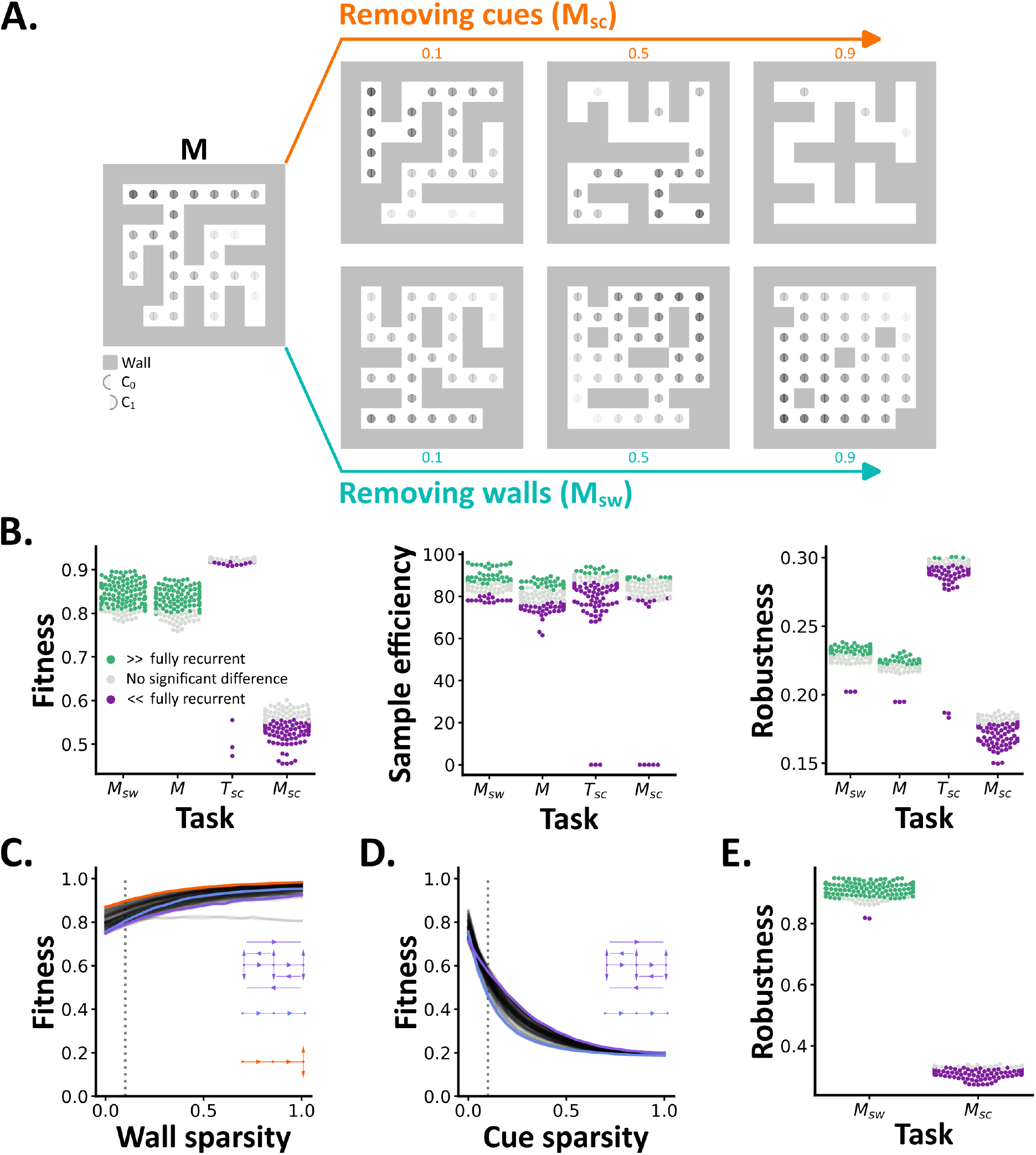
**A** Multisensory mazes. We create maze structures, shown with walls in grey and paths in white, and fill these with gradients of sensory cues (circles). We introduce further complexity by randomly removing some fraction of walls or cues. Seven maze structures are shown with various levels of wall and cue sparsity. **B** The median fitness (left), sample efficiency (middle) and robustness to sensor noise (right) across 50 trained networks, per task, for all 128 architectures (circles). Architectures which perform significantly better or worse (*p <* 0.01) than the fully recurrent architecture are shown in green and purple, the rest are shown in grey. **C-D** Each architecture’s median fitness as a function of increasing wall (C) or cue sparsity (D). All architectures are drawn in grey, with select architectures highlighted in colour. Dashed vertical lines denote the level of sparsity during training. Note that here, feedforward denotes the large-feedforward architecture. **E** Statistical comparisons, as in B, comparing each architecture’s robustness to wall or cue sparsity to the fully recurrent architecture.

When navigating in environments with dense sensory cues (M_sw_ and M), every pRNN architecture achieved equivalent or significantly better fitness than the fully recurrent architecture (Figure 3B, left); with the best achieving 11% and 9% higher fitness. Moreover, 30% of architectures were significantly more sample efficient (Figure 3B, middle) and 47% were significantly more robust to noise (Figure 3B, right). Conversely, when navigating in environments with sparse sensory cues (T_sc_ and M_sc_), on average across all three metrics, 41% of architectures performed worse than the fully recurrent architecture, and the rest performed equivalently (Figure 3B). Additional experiments with homogenous activation functions per layer, and smaller and larger networks, further support these findings (section 6.3).

Together, these results demonstrate that partially recurrent networks can outperform fully recurrent networks in environments with dense sensory cues, and perform equivalently in more complex scenarios. Though, perhaps the main advantage of fully recurrent networks is that they are better able to adapt, or generalise, to novel situations?

To explore this, we tested how well networks trained at one level of wall (M_sw_) or cue sparsity (M_sc_) performed when tested across a wider range. In general, fitness increased as a function of wall sparsity (Figure 3C) - as the average path length from start to goal decreased, but decreased with cue sparsity (Figure 3D) – as the number of informative cues decreased. Most pRNN architectures generalised significantly better to wall sparsity than the fully recurrent architecture, while the opposite was true for cue sparsity (Figure 3E); illustrating that fully recurrent networks do not necessarily generalise better.

In sum, these results demonstrate that partial recurrence enables efficient, yet performant solutions. However, how does structure link to function across these architectures and tasks? For instance, in the track maze task the architecture which is most robust to noise, includes all three backwards pathways and the most sample efficient architecture includes forward skip connections (Figure 2B). Do these examples highlight general structure-function principles?

### 3.5 Each pathway’s functional impact is circuit and task dependent

To explore the link between network structure and function, we used counterfactual analysis (Verma et al., 2021; Guidotti, 2024) to ask: do pairs of architectures, which differ by only a single pathway, differ significantly in their function? The architectures in fig. 4B (left) are one such pair, as they are identical, except for the backwards connection *W*_*hi*_ – present in one and absent in the other.

**Figure 4:**
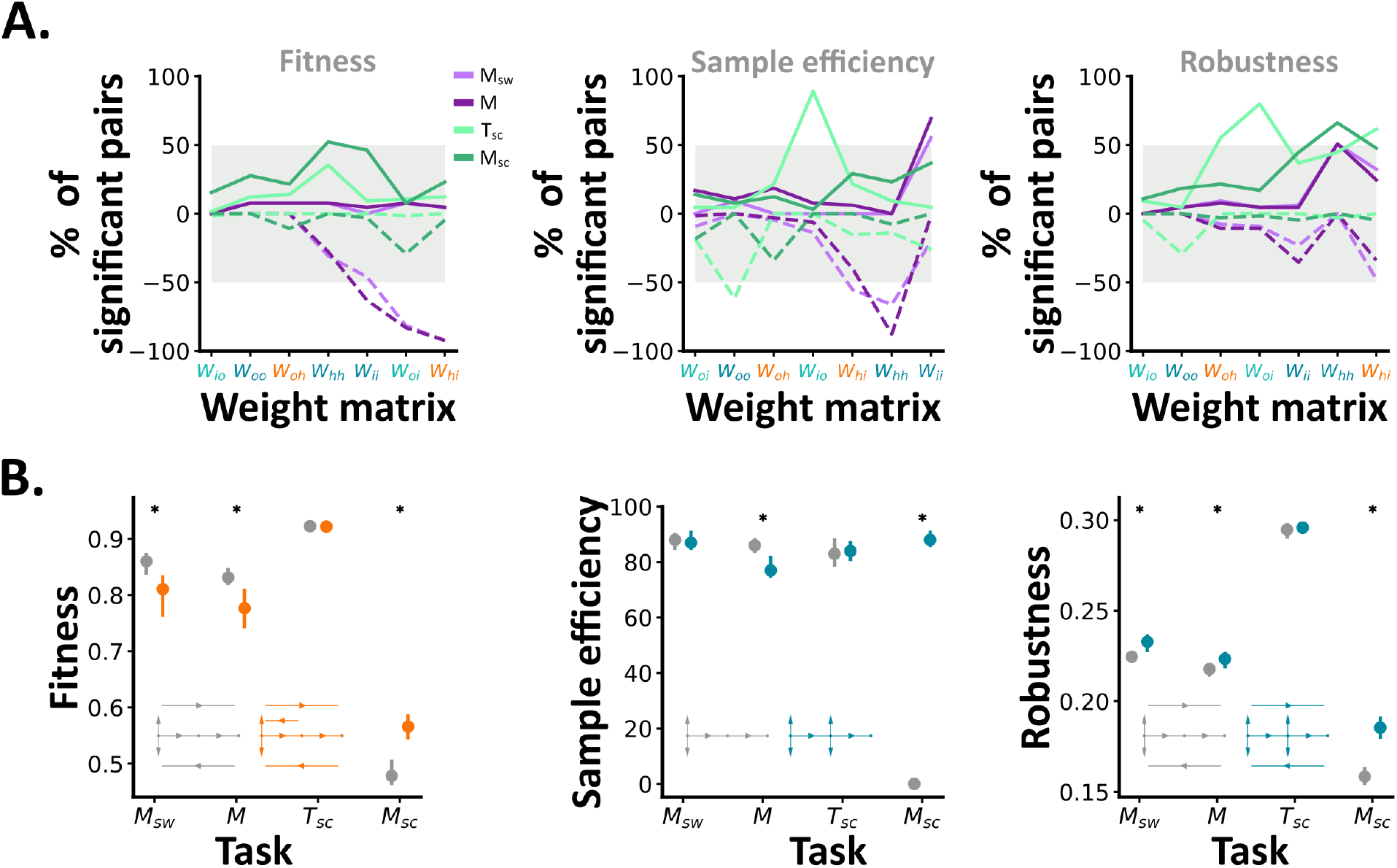
**A** The percentage of counterfactual pairs in which the architecture with an additional weight matrix has significantly greater (solid lines) or less (dashed lines): fitness (left), sample efficiency (middle) or robustness to sensor noise (right). Results from each task are shown in distinct colours. The grey underlay marks the region in which most differences are context dependent. **B** Example counterfactual pairs with task dependent differences in fitness (left), sample efficiency (middle) and robustness to sensor noise (right). For each pair we plot the two architectures, and the median and interquartile range for each task. Statistically significant differences are marked with a * (*p <* 0.01).

Specifically, for a given measure of function *y* and a pair of architectures which differ by one pathway *i*, we test if the distribution *y*_*i*=1_ (across 50 networks) is stochastically less than or greater than *y*_*i*=0_ (another set of 50 trained networks). From this approach we draw two main conclusions.

First, each pathway’s functional impact is *circuit dependent*. If a pathway had a consistent effect on function, for example increasing robustness to sensor noise, then all or most networks with that pathway would differ significantly from their counterfactual pairs. Instead, for almost all pathways and functional metrics, we observe significant differences in less than 50% of pairs (Figure 4A). For example, on average across tasks, only 42% of networks with backwards hidden-input (*W*_*hi*_) connections are significantly more robust to sensor noise than their pairs without (Figure 4A, right panel, rightmost x tick, solid lines).

Second, each pathways functional impact is *task dependent*. For instance, when trained and tested in mazes with dense sensory cues (M_sw_ and M) 92% of networks with additional hidden-input connections (*W*_*hi*_) were significantly less fit than their pair, conversely in mazes with sparse cues (T_sc_ and M_sc_) only 2% were less fit (Figure 4A, left, purple vs green dashed lines). These task dependent effects were even evident at the level of single circuit pairs. For example, the architecture with *W*_*hi*_ connections in fig. 4B (left panel, orange) was significantly less fit than it’s pair (grey) in dense mazes, yet performed equivalently or better in sparse mazes. We observed similar effects for both sample efficiency and robustness to sensor noise (Figure 4B, middle and right).

Overall, these data demonstrate that, depending on the circuit and task, small changes to network structure can lead to dramatic changes in function. Though, *why* do these changes alter network function?

### 3.6 From structure to function via computational traits

To explore why changes to network structure alter network function, we measured three computational traits from every trained network: their pathway magnitudes, input-output sensitivity and memory dynamics. Then, we quantified how predictive these traits are of network function. For example, given a network’s memory dynamics, how accurately can one predict its robustness to noise? These results lead us to propose a conceptual model, whereby: networks with different structures learn distinct computational traits, which determine their function.

To describe each network’s pathways we calculated the magnitude of each weight matrix (∥*W*_*i*_∥ _*F*_ ). These values were surprisingly consistent across architectures and tasks: with large feedforward and forward skip magnitudes, and relatively low values otherwise (Figure 5A).

**Figure 5:**
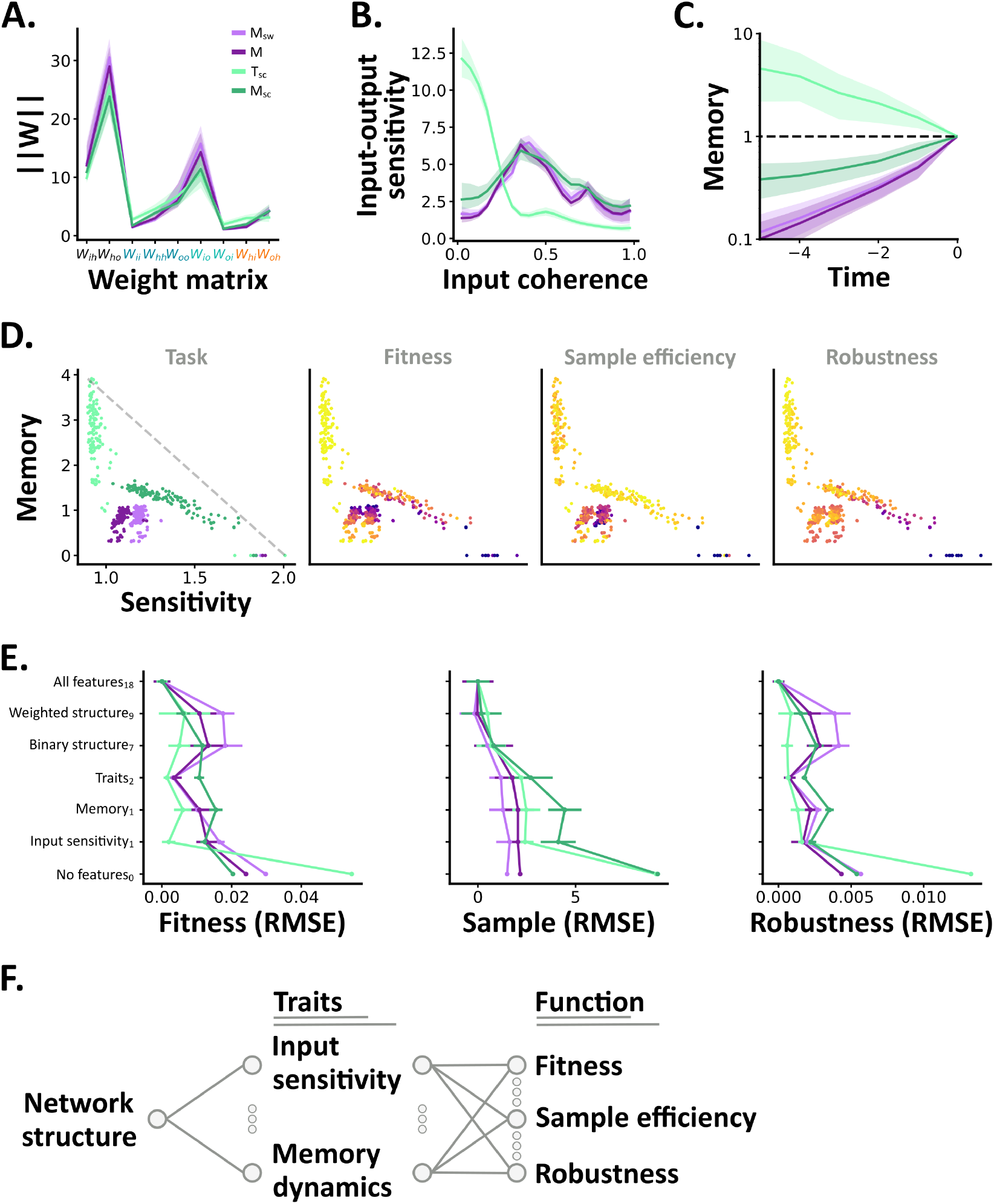
**A** The learned magnitude of each weight matrix. Results from each task are shown as median (solid line) plus interquartile range (shaded surround) across all trained networks. **B** Input-output sensitivity as a function of input coherence plotted as in A. **C** Memory as a function of time plotted as in A. **D** Input sensitivity vs memory (log-log plots). Each circle represents the median of a single architecture from 50 trained networks per task. In the left panel architectures are coloured by the task they are trained on. The dashed grey line illustrates a trade-off between input sensitivity and memory – simply plotted from: (min(x), max(y)) to (max(x), min(y)). In the other subplots circles are coloured by their fitness, sample efficiency and robustness from low (blue) to high (yellow) within each task. **E** Random forest models trained to predict fitness (left), sample efficiency (middle) and robustness to sensor noise (right) using different computational traits; subscripts note the number of input features per model. We plot the mean error plus standard deviation from 10-fold cross-validation, with individual lines per task. **F** A conceptual model showing how networks with different structures learn solutions with distinct computational traits which influence their function. The small circles, between labelled nodes, suggest additional computational traits or functional metrics.

To measure each networks input-output sensitivity and memory dynamics, we collected data from 1, 000 test trials per network and defined a metric to describe how sensitive each network’s outputs *A*, at time *t*, are to its inputs *C* at time *t* − *k* (*k* ≥ 0). Where the elements of *C* represent the inputs from each sensor, and the elements of *A* represent the network’s output for each possible action.

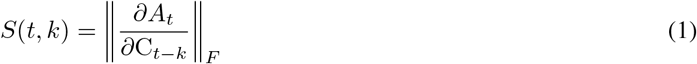

We define a network’s *input-output sensitivity* as the mean ⟨·⟩ of its sensitivity at time *t* to inputs at time *t*: ⟨*S* = *S*(*t*, 0) ⟩ . Calculating this metric from networks trained on the track maze task (T_sc_), revealed an intuitive result: when the inputs to a network clearly indicate one direction (e.g. left *>* right), input sensitivity is low. And so, networks are relatively insensitive to changes in their inputs. By contrast, when inputs are ambiguous (i.e left ≈ right) networks are highly sensitive to changes in their inputs (Figure 5B, light green line). Networks trained on our other tasks showed a different sensitivity profile, with low sensitivities at both low and high levels of input coherence, and high values in-between (Figure 5B, dark green and purple lines). This may suggest that, in these tasks, networks learn to: filter out ambiguous inputs, detect subtle differences in semi-ambiguous inputs and simply follow clear inputs. Though, this is just one possibility and individual architectures may leverage different strategies.

We define *memory at lag k* to be the ratio of the sensitivity to past compared to current inputs, *M*_*k*_ = ⟨*S*(*t, k*)*/S*(*t*, 0) ⟩ . As such, as *M*_*k*_ increases, networks transition from being completely insensitive to their prior inputs (when *M*_*k*_ = 0) to being more sensitive to their prior than current inputs (when *M*_*k*_ *>* 1). In the track maze task (T_sc_), we found that most networks’ memory increased further back in time (Figure 5C, light green line), meaning that their outputs at any time, are more sensitive to changes in the past than the present. This approach works for this maze design as, assuming an agent sets off in the correct direction, repeating its first action will take it to the goal.

However, in more complex scenarios (M_sw_, M, M_sc_), most networks’ memories decayed with time (Figure 5C, dark green and purple lines) showing that they are more sensitive to recent than past inputs; though the rate of this decay, was task-dependent. Specifically, networks trained on the maze with sparse cues (M_sc_) were more sensitive to their past inputs than networks trained on the tasks with dense cues (Figure 5C dark green vs purple lines).

To explore the link between these computational traits and network function, we began by visualising the relation between each architecture’s mean input sensitivity, total memory and functional measures (Figure 5D). This showed that different architectures learn distinct input-output sensitivities and memory dynamics, and revealed a trade-off between the two; where improving one impairs the other. This is likely due to the fact that retaining hidden states improves memory, but combining those hidden states with current inputs interferes with the direct mapping of inputs to outputs.

Finally, to quantify the relation between these traits and our functional measures, we used random forest regression models – which can capture complex, nonlinear relations between different combinations of features and continuous values. As a lower bound, we calculated the error from simply predicting the mean of each functional measure. As an approximate upper bound, we fit random forest models to all 18 of our features of interest.

Random forest models trained to predict function from each network’s weighted structure, performed similarly to models trained on binary network structures (Figure 5E) despite having an additional two parameters, and using continuous instead of binary values. This demonstrates that knowing each pathways magnitude does not provide much additional information over simply knowing if it is present or absent. By contrast, both input sensitivity and memory were highly predictive of function, achieving errors far lower than guessing the mean and close to or even better than models with many more features (Figure 5E). For instance random forest models trained to predict robustness to sensor noise from memory achieved a mean normalized error of 0.002 across tasks; better than our lower bound of 0.007 and as good as models trained on each network’s weighted structure (0.002) – despite using ^1^*/*_9_ the number of features (Figure 5E, right).

All together, these results lead us to propose that networks with different structures learn distinct input-output sensitivities and memory dynamics, which in turn influence their function (Figure 5F). This model is primarily supported by two pieces of evidence. First, across architectures we observe a diverse range of input-output sensitivities and memory dynamics (Figure 5D). Second, knowing these traits allows us to predict network function with high accuracy (Figure 5E). As these two traits describe how networks process and retain information, they are likely to be relevant to many tasks, and would be interesting to approximate from real neural circuits. More broadly, we hope that providing a conceptual stepping stone, between structure and function, will help to bridge this gap.

## 4 Discussion

Many neural circuits sit somewhere between feedforward and fully recurrent. How does this structure shape their function? To explore this question, we define: a neural network model (pRNNs), a new set of tasks (multisensory mazes) and tools for linking network structure to function via computation. Below, we discuss our findings in the context of neuroscience and machine learning.

In biological circuits, connections cost both energy and space. So, *a priori* partially recurrent networks should be more efficient than fully recurrent networks. However, does this efficiency come at the cost of performance?

While prior studies have found that sparsity comes with at least a small cost to performance in task-trained models (Bellec et al., 2017; Jayakumar et al., 2020; Goulas et al., 2021; Achterberg et al., 2023; Béna and Goodman, 2025), other work has shown that sparse architectures can support robust associative memory and efficient computation (Hoffmann, 2019). Our findings reconcile these perspectives by demonstrating that partially recurrent networks — which are both sparse and bidirectional — can achieve high performance, sample efficiency, and robustness, depending on their structure and the task. We attribute this difference to two aspects of our approach.

First, our choice of tasks. In general prior studies have focussed on cognitive neuroscience or machine-learning style tasks in which networks must generate a single response to a sequence of inputs (Goulas et al., 2021; Achterberg et al., 2023; Liu et al., 2024; Béna and Goodman, 2025). By contrast, our maze tasks require networks to continuously translate the inputs from noisy sensors (and internal states), into actions; while being simple enough for us to focus on relatively small neural networks (with between 104 and 442 learnable parameters) and to train an exceptionally large number of networks (50 per architecture and task → over 25,000 trained networks). Working at this level of task complexity, resource constraints and statistical power, allowed us to observe differences between classical models of multisensory integration (section 3.2), and between our partially recurrent architectures (section 3.4). We therefore anticipate that these tasks will serve as a valuable foundation for future studies.

Second, our choice of model. Many prior studies have varied network structure by taking recurrent neural networks and constraining their hidden-hidden weight matrix (*W*_*hh*_) to generate networks with distinct properties. However, within our framework, these networks are all variations on a single architecture; and so, by comparing every pRNN architecture we explored a far more diverse set of network structures.

That said, one limitation of our study is that we only consider models with a single hidden layer. Future work should aim to explore deeper models, closer to biological networks in both their structure and function. However, the number of pRNN architectures scales super-exponentially with the number of layers (L): 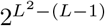. As such, complete searches of the space, as we present here, quickly become infeasible. One solution to this would be to develop search algorithms to navigate the space of pRNN architectures – akin to how Castro et al. (2025) explored the space of symbolic cognitive models. However, our results demonstrate that even relatively small changes to network structure can lead to dramatic changes in function (section 3.5); suggesting that this may be challenging. Another solution would be to use neuroevolution algorithms (Stanley and Miikkulainen, 2002; Stanley et al., 2019). These algorithms optimise a given function (e.g. task performance) by iteratively evaluating and varying a population of neural network architectures. The advantage of this approach is that it enables the discovery of diverse solutions (Gaier and Ha, 2019). The downside is that it can be hard to interpret large collections of neural networks.

In this direction, we measured three computational traits from all of our trained networks. We found that different architectures learned distinct input-output sensitives and memory dynamics, and observed a trade-off between the two (section 3.6). As these metrics essentially quantify how networks how networks process and retain information, they are likely to be relevant for many tasks in both neuroscience and machine learning, and could even be approximated from real circuits via connectome-based models (Lappalainen et al., 2024; Shiu et al., 2024). Though, other computational traits may be more relevant for different tasks or architectures, and developing these remains an exciting direction for future work.

Finally, we demonstrate that our computational traits are highly predictive of function. For instance, knowing a network’s memory dynamics allows us to accurately predict its robustness to noise (section 3.6). These results lead us to propose that networks with distinct architectures learn solutions with different computational traits which influence their function. This conceptual model suggests a shift in focus – from identifying *which* network structures yield different functions, to understanding *why* they do so, via their computational traits. And sparks numerous questions: to what extent do computational traits explain the functional differences between other neural network architectures? Could optimising networks for specific traits, rather than specific tasks, yield more flexible solutions? And could traits like these be key to understanding inter-circuit, inter-individual or even inter-species differences? Ultimately, our work provides a springboard for addressing fundamental questions, like these, related to how structure sculpts function in artificial and biological networks.

## 5 Methods

In brief, we compare how agents with different policies perform on a set of maze tasks (section 5.2). These tasks are set in grid environments, with impassable walls, paths the agents can move on, and sensory cues which indicate the shortest path through each maze. To perform these tasks, each agent implements a three-step sensation-action loop (section 5.3). First, they sense their local environment. Often, we add independent Gaussian noise to each sensor at each time step. Next, they implement a policy – either an algorithm (section 5.4) or a forward pass through a neural network (section 5.5). Finally, agents choose and implement an action from a discrete set; either pausing or moving in a cardinal direction. To compare models we measure three functional metrics (section 5.6) and three computational traits (section 5.7). For statistical comparisons across models we use Mann–Whitney U tests with corrections for multiple comparisons (section 5.8).

### 5.1 Code

All code was written in Python and is available, as a fully documented Pip installable package, at https://github.com/ghoshm/Multimodal_mazes. All results were obtained from a single version of the code (Git tag: multimodal_mazes-v1). All trained models, and aggregated results, will be deposited in Zenodo upon publication.

### 5.2 Multisensory mazes

All tasks are set on discrete grids, with tracks the agents can move on, and impassable walls. All tasks are run in discrete time, for a fixed number of time steps (see section 5.9.1 for task hyperparameters).

#### 5.2.1 Track maze

In this task, agents start in the middle of a track (Figure 1C, left). Their aim is to reach a goal location at either the left or right end of the track within a fixed number of time steps. To do so, they must use sensory cues from two channels (C_0_ and C_1_). In each trial, one channel’s cues are linearly spaced between 0 and 0.5, leading from start to goal. While the other channel’s cues are linearly spaced between 0 and 0.5 from start to both ends of the track, and so are ambiguous. Across trials we vary both the goal location and whether each channel is leading or ambiguous, yielding 4 configurations. Across configurations, the optimal strategy is to sum the information over both channels and head in the direction with the greatest total. For example, at the first time step, with the goal on the left, and no sensor noise, an agent may sense:

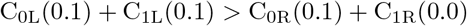

We also consider two variants of this task.

**Nonlinear track maze** - we double the strength of the cues leading away from the goal (Figure 1C, middle). This ensures that, at the first time step, there is an equal amount of information either side of the agent and a linear strategy is uninformative. Following the example above, the agent would instead sense:

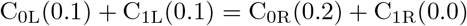

Now the optimal strategy is to use a nonlinear function to weight the multisensory signal (leading towards the goal) more strongly than the unisensory signal (leading away from the goal).

**Track maze with a gap** - we introduce a gap in the signals leading towards the goal, by setting both channels to zero *m* steps prior to the goal (Figure 1C, right). I.e. when *m* = 1, the location prior to the goal has no sensory information, when *m* = 2 the two locations prior to the goal have no information. Consequently, the optimal strategy is to set off in the direction with the greatest total signal, at the first time step, and to persist in that direction.

#### 5.2.2 General mazes

In these tasks, agents start in one corner of a maze (set on an *n × n* grid) and must navigate to a goal location in another corner (Figure 3A). To generate each maze’s structure (walls and paths) we use the Aldous-Broder algorithm (Aldous, 1990; Broder, 1989). We then choose random corners for the start and goal locations, and fill sensory cues into each channel, based on the shortest-path distance from each location to the goal. These cues are linearly spaced between: 0 - the furthest point from the goal (which is not necessarily the start position), to a maximum of 1*/c* at the goal location, where *c* is the number of sensory channels. Then, we introduce additional complexity by varying two parameters.

**Wall sparsity** - prior to filling in the sensory cues, we randomly remove some fraction of walls (excluding the maze’s outer wall). Increasing this parameter from 0 to 1 interpolates between a perfect maze and an open field.

**Cue sparsity** - after filling in the sensory cues, we randomly set some fraction of cues (in both channels) to 0. Increasing this parameter from 0 to 1 interpolates between a maze with sensory cues at every location to a maze with no sensory cues.

### 5.3 Agents

Each agent occupies a location on a discrete grid, and implements a three-step sensation-action loop (Figure 1A):

1. **Sense** - each of the agent’s sensors reads one channel’s signal from one adjacent square (e.g. C_0L_, C_1L_, C_0R_ etc). Independent Gaussian noise is optionally added to each sensor at each time step. If the agent is at (*x*_*a*_, *y*_*a*_), then the reading from channel *i* in direction *j* is given by C_*ij*_ = Cue(*i, x*_*a*_ + Δ*x*_*j*_, *y*_*a*_ + Δ*y*_*j*_) + 𝒩 (0, *σ*^2^), where *σ* = 0 in the noise-free case. Sensor values are clipped to the range [0, 1].
2. **Policy** - maps sensor inputs to an output vector with one value for each possible action. Policies are either multisensory algorithms (section 5.4) or neural networks (section 5.5). Some policies are stateless, i.e.. they map sensations to actions independently at every time step, but others combine their sensory inputs with internal (hidden) states.
3. **Act** - choose and implement an action: using argmax(*A*) where *A* represents a vector of potential actions (either pausing or moving in a cardinal direction). When multiple actions are equally likely, agent’s choose randomly between them. When an agent tries to move into a wall, it instead remains in place for that time step.

### 5.4 Multisensory algorithms

Throughout we use the term *algorithm* to describe policies which map a vector of sensory inputs (C) to a vector of potential actions (*A*), without using any learnable parameters or hidden states. C represents the agent’s sensors (left, right, up, down), where C_ij_ denotes the values of sensory channel *i* in direction *j. A* represents possible actions (move left, right, up down), where *A*_*j*_ is the value of the action in direction *j*. At each time step agents choose argmax(*A*) (with randomization between equal values). We set *A*_*j*_ = 0 for all *j*, at the start of every time step.

We explore the following 7 algorithms (Figure 1B).

#### 5.4.1 Random motion

Choose randomly between all actions:

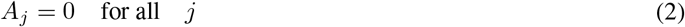

#### 5.4.2 Unisensory

Map each sensors value, from channel *i*, directly to the corresponding output action:

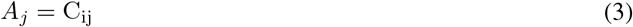

#### 5.4.3 2-look binary

Within each channel identify the sensor with the greatest activation *d*(*i*). If they agree, choose the corresponding action. Otherwise, choose randomly between the suggested actions:

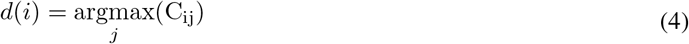

For each *i* update:

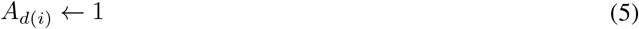

#### 5.4.4 2-look max

Within each channel identify the sensor with the maximum activation *d*(*i*). Add the activations from these sensors to their corresponding outputs.

For each *i* update:

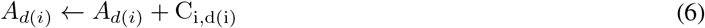

#### 5.4.5 Max-dv

Map the maximum value from each sensor pair (e.g. the left sensors) to the corresponding actions:

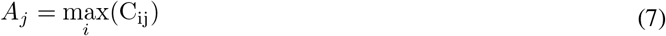

#### 5.4.6 Linear fusion

Sum the sensor signals across channels:

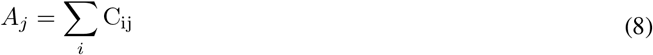

#### 5.4.7 Nonlinear fusion

Sum the sensor signals across channels and add a nonlinear term. Here, we use the square root of their product, but any nonlinearity would suffice (Ghosh et al., 2024):

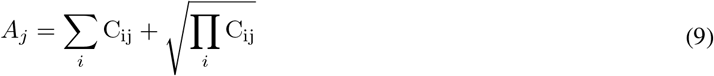

### 5.5 Partially recurrent neural networks

#### 5.5.1 Model

Given a neural network with: an input, a hidden and output layer, there are 9 possible weight matrices:

- **Feedforward** connections - from the input to hidden layer (*W*_*ih*_) and hidden to output layer (*W*_*ho*_).
- **Lateral** connections - between the units within each layer: input-input (*W*_*ii*_), hidden-hidden (*W*_*hh*_) and output-output (*W*_*oo*_). Note that these include self (unit_*b*_–unit_*b*_) connections.
- **Skip** connections - from the input to output layer (*W*_*io*_) and output to input layer (*W*_*oi*_).
- **Backwards** connections - from the hidden to input layer (*W*_*hi*_) and output to hidden layer (*W*_*oh*_).

Assuming the feedforward connections (*W*_*ih*_ and *W*_*ho*_) are always present, and each of the other 7 matrices can be present in any combination yields 2^7^ = 128 architectures. Relaxing this assumption would generate 2^9^ = 512 architectures, though some of these would have no path from input to output. When none of these 7 additional matrices are present the model is purely feedforward. When all of the additional matrices are present the model is fully recurrent as every unit connects to every other unit bidirectionally. We term the remaining 126 architectures, between these two extremes, *partially recurrent neural networks*.

Units in the input, hidden and output layers (respectively) use linear, rectified linear (ReLU) and softmax activation functions, with no bias terms.

In the feedforward case we write the forward pass (or policy *π*) as:

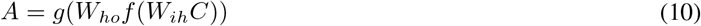

Where *A* represents a vector of potential actions (move left, right, up, down or pause). *C* represents the agent’s sensors (left, right, up, down from two channels) *W*_*ih*_ and *W*_*ho*_ are the two feedforward weight matrices, *f* is ReLU and *g* a softmax across possible actions.

In the fully recurrent case, the forward pass (or policy *π*) is:

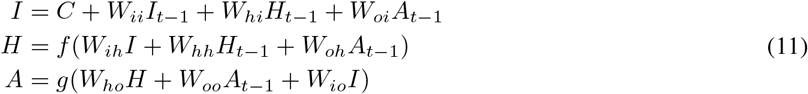

Where *I* and *H* represent the activations of the input and hidden units at time *t*. And all hidden states are set to zero at the start of each trial.

In partially recurrent cases, each model’s forward pass follows Equation (11) with any absent weight matrices (determined by the model’s structure) set to zero. See section 5.9.2 for model hyperparameters.

#### 5.5.2 Training

To optimise each model’s weights we used deep Q-learning (Mnih et al., 2013; Jensen, 2024). For each time step within each training episode, each agent senses its environment, chooses an action and acts. We use an epsilon-greedy action selection policy, where agents initially explore by choosing actions at random, but increasingly rely on their learned policy *π* as training progresses. Specifically, we define a vector ***ϵ*** with one value per training maze, which decreases step-wise from 0.95 to 0.25 over training.

At each time step, the agent selects an action *a*_*t*_ according to:

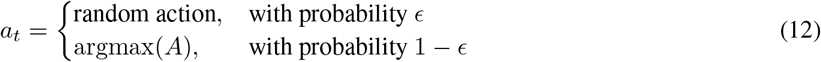

At each time step, agent’s estimate the value of the action they took (termed the Q-value) using their policy (a forward pass through a neural network):]

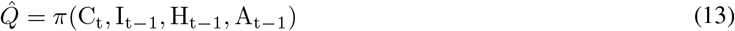

We calculate the target Q-values:

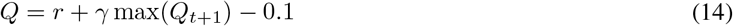

Where the reward *r* is proportional to the reduction in shortest-path distance to the goal, scaled by a factor of 2. *γ* is a discount factor that determines how strongly the agent values future rewards. max(*Q*_*t*+1_) is the maximum predicted Q-value at the agent’s new location, and the constant -0.1 introduces a small penalty for each time step to encourage faster solutions.

Next, we calculate the mean squared error between the two (the loss):

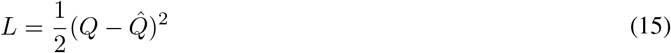

We accumulate these losses, then at the end of every episode update the agent’s weights using backpropagation. Episodes terminate after a fixed number of time steps or the goal being reached. As each model’s forward pass is written in PyTorch (Paszke et al., 2019), we rely on autodifferentiation for each model’s backward pass.

### 5.6 Functional metrics

To describe each model’s behaviour, we define three functional metrics.

#### 5.6.1 Fitness

To measure performance in our maze tasks we define the following fitness function:

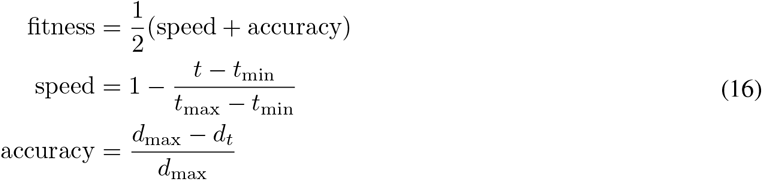

Where *t* is the time taken to reach the goal, *t*_min_ is the quickest time possible, *t*_max_ is the maximum number of steps in the trial, *d*_max_ is the furthest distance from the goal and *d*_*t*_ is the agent’s distance from the goal at time *t*. Note that when the goal is not reached *t* = *t*_max_. This fitness function ranges from 0 - ending the trial at the furthest point from the goal, to 1 – taking the shortest path from start to goal.

We define each model’s fitness by testing it on 1,000 mazes. For our neural network models we do this after training with all parameters set as they were during training (i.e. in distribution).

#### 5.6.2 Sample efficiency

To measure a network’s learning speed, we record it’s test fitness 100 times, at equally spaced intervals, throughout training. This generates a vector, per network, *F*_*t*_ - the test fitness after *t* epochs. We define sample efficiency as 100 minus the point at which: *F*_*t*_ is first 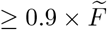 where 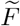 denotes the median fitness across the entire set of networks trained on a given task. As such, higher values represent better sample efficiency. Networks which do not reach this threshold are assigned a value of 0.

#### 5.6.3 Robustness

To test an agent’s robustness, we repeatedly test it on a set of 1,000 mazes as we: increase its sensor noise (e.g. Figure 1D) or remove an increasing fraction of walls (Figure 3C) and sensory cues (Figure 3D) from each maze. We test: 21 values of noise (linearly spaced between 0 and 0.5) and 21 values of wall and cue sparsity (linearly spaced between 0 and 1). For our pRNNs we test robustness after training, with all other parameters kept in distribution. To summarise each agent’s robustness we compute the area under each curve (fitness vs noise or sparsity) using the trapezoidal rule.

### 5.7 Computational traits

To explore why changes to network structure alter network function, we measure three computational traits from every network after training.

#### 5.7.1 Pathway magnitudes

For each network we calculate the Frobenius norm 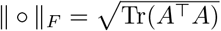 of each weight matrix ∥*W*_*i*_∥_*F*_ (Figure 5A).

#### 5.7.2 Input-output sensitivity

Following training we test each network on a set of 1,000 mazes, with all parameters set in distribution. At every time step during these trials we collect each network’s inputs (*C*), and the states of their input (*I*), hidden (*H*) and output (A) units. For each input state we run a forward pass through the trained network. Then, following Equation (1), we calculate the input-output Jacobian and take the Frobenius norm ( ∥ ∘ ∥_*F*_ ); using the Jacobian and norm functions from PyTorch (Paszke et al., 2019).

When input-output sensitivity is high, small changes in the network’s inputs will lead to large changes in its outputs. Conversely, when input-output sensitivity is low, changes to the network’s inputs affect the outputs less. To obtain a single input-output sensitivity value per network we calculate the mean norm across all states.

To plot each network’s input sensitivity as a function of input coherence (*c*) as in fig. 5B, we let:

- 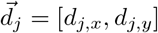 be the unit direction vector for direction *j* ∈ {left, right, up, down},
- *m*_*j*_ = ∑_*i*_ C_*ij*_ be the total magnitude of sensory input in direction *j* across all channels *i*,
- 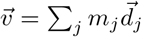 be the net input vector,
- *M* = ∑_*j*_ *m*_*j*_ be the total input magnitude.

Then we define input coherence *c* as:

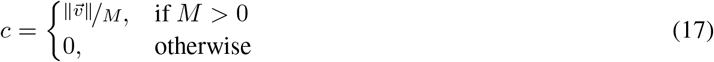

where 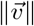 denotes the Euclidean norm of the vector 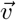, which is always between 0 and 1. When *c* is close to 1, the input state clearly indicates the goal direction, while when *c* is low the inputs are ambiguous.

#### 5.7.3 Memory dynamics

Following training we test each network on 1,000 test trials, with all parameters held in distribution, and collect all of the network’s states at every time step. Following Equation (1), we define a network’s memory at temporal lag *k* (*k* ≥ 0) as: *M*_*k*_ = ⟨*S*(*t, k*)*/S*(*t*, 0) ⟩ . Where, *S*(*t, k*) represents the sensitivity of the network’s outputs at time *t* w.r.t. its inputs at time *k, S*(*t*, 0) its sensitivity to its current inputs, and ⟨·⟩ denotes the mean across all input states. As such, when *M*_*k*_ = 0 a change in the network’s inputs at lag *k*, will have no effect at time *t* (as in a stateless model). When *M*_*k*_ *>* 1, networks are more sensitive to their prior than current inputs. And, when 0 *< M*_*k*_ *<* 1, networks are sensitive to their prior inputs, but more sensitive to their current inputs. To obtain a single memory value per network we sum each network’s memory across all temporal lags. When drawing comparisons between networks trained on tasks with different trial lengths, as in fig. 5D, we only sum lags up to the shortest trial length.

### 5.8 Analysis

#### 5.8.1 Statistical comparisons

For a given metric, e.g. fitness, we compare each architecture (a distribution across 50 networks) to the fully recurrent architecture (a second distribution across 50 networks), using two-sided, asymptotic Mann–Whitney U tests. We then adjust the resulting (129) p-values, using the Benjamini-Hochberg procedure, to control the false discovery rate at

0.01. For instance, if 100 architectures are found to differ significantly from the fully recurrent architecture after correction, we would expect approximately 1 of these to be a false positive. Practically we used the `mannwhitneyu` and `false_discovery_control` functions from SciPy (Virtanen et al., 2020).

#### 5.8.2 Random forest models

To quantify the relation between our computational traits and our functional measures, we used random forest regression models. Starting from a binary matrix *X* with a row per trained network and a column per feature, and a vector *y* with a value per trained network. We fit random forest models (*f*), using different subsets of features (*X*_*i*_): ŷ = *f*(*X*_*i*_). Practically we used the `RandomForestRegressor` function from scikit-learn (Pedregosa et al., 2011) with 10-fold cross validation. To mitigate overfitting, we limited the depth of each tree to a maximum of 9 levels using the `max_depth` parameter.

### 5.9 Hyperparameters

#### 5.9.1 Task hyperparameters

For our 4 tasks we used the following hyperparameters.

**Table 1:** A table noting the hyperparameters we used for each task: track mazes with sensory gaps one step prior to the goal (T_sc_), 2D mazes with dense walls and sensory cues (M), 2D mazes with 10% of walls removed (M_sw_), and 2D mazes with 10% of sensory cues removed (M_sc_). # denotes the number of mazes used for training and testing.

| Task | Size ($n \times n$) | Trial length | # Training | # Testing | Wall sparsity | Cue sparsity |
| --- | --- | --- | --- | --- | --- | --- |
| T<sub>sc</sub> | 11 | 6 | 200,000 | 1,000 | 0.0 | 1 gap |
| M | 9 | 20 | 400,000 | 1,000 | 0.0 | 0.0 |
| M<sub>sw</sub> | 9 | 20 | 400,000 | 1,000 | 0.1 | 0.0 |
| M<sub>sc</sub> | 9 | 20 | 400,000 | 1,000 | 0.0 | 0.1 |

#### 5.9.2 Model hyperparameters

We used the following hyperparameters for all of our neural network models. Except for the large feedforward model which used 34 hidden units - to match the fully recurrent architecture’s number of trainable parameters. Results from networks with homogenous activation functions per layer, and smaller and larger networks are shown in section 6.3.

**Table 2:** A table listing the hyperparameters we used for our neural network models. Hyperparameters only used during training are marked with a *. We used Epsilon-greedy action selection, with a linear, stepped decay schedule from 0.95 to 0.25. We used the `clip_grad_norm` function from PyTorch (Paszke et al., 2019) to clip our gradients to a maximum value of 10.

| Hyperparameters | Value |
| --- | --- |
| Input units (#) | 8 |
| Hidden units (#) | 8 |
| Output units (#) | 5 |
| Sensor noise scale | 0.05 |
| Learning rate* | 0.001 |
| Gamma* | 0.9 |
| Epsilon decay schedule* | see caption |
| Movement penalty* | 0.1 |
| Gradient clipping* | see caption |

## 6 Supplementary information

### 6.1 Network rank does not directly relate to structure

Some studies vary the rank of a recurrent neural network’s hidden-hidden weight matrix and consider this analogous to changing its structure. However, consider a network with a three nodes, layers or areas and the following, weighted connectivity matrix:

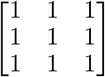

This matrix is fully recurrent and is of rank 1. Now, if we double the strength/weight of the first connection

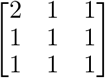

the rank becomes 2. While this will change how signals propagate through the network/graph, it’s structure is still fully recurrent.

Moreover, if we consider all 128 of our architectures as binary 3 *×* 3 matrices (representing each architectures: input, hidden and output layers and the presence or absence of connections between them), then we find that these diverse network structures are all of rank 1, 2 or 3. This further demonstrates that rank does not uniquely determine or reflect network structure.

### 6.2 Architectures

**Figure 6:**
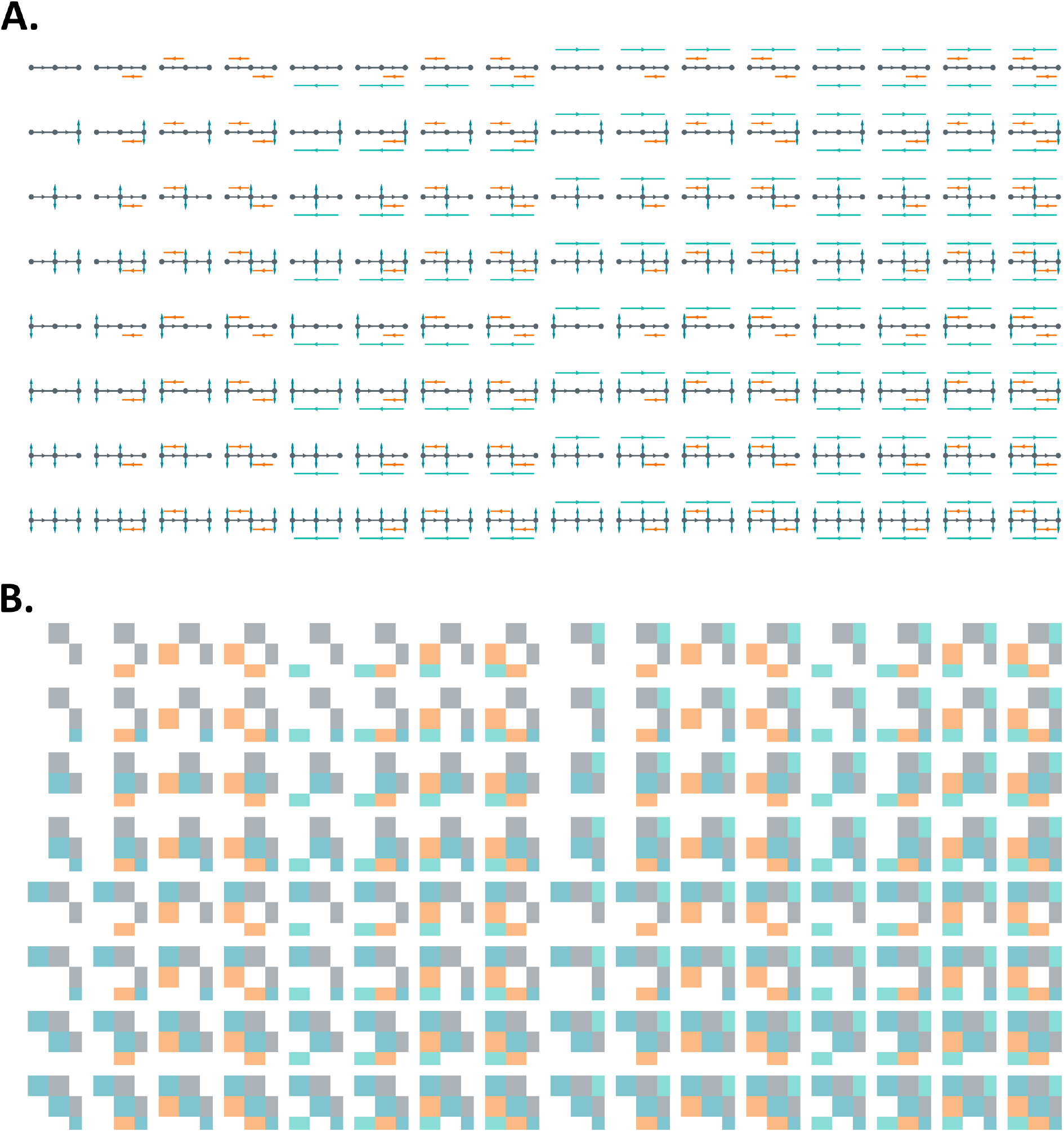
**A** All 128 architectures drawn as pathway models. Each model is composed of: input, hidden and output layers (represented by grey circles), two feedforward pathways (grey lines) and any combination of 7 other pathways. Lateral pathways are drawn in blue, skips in green and backwards in orange. **B** All 128 architectures drawn as sparse unit-unit matrices. Each subplot shows each architectures full unit-unit weight matrix (including input, hidden and output units). Feedforward weights are drawn in grey, lateral in blue, skip in green and feedback in orange. White indicates absent connections.

### 6.3 Hyperparameter robustness

To test the robustness of our results to different hyperparameter choices, we trained four additional sets of networks. For each set, we trained all 128 architectures (with 50 repeats per architecture) on our hardest task (the maze with sparse cues, M_sc_).

Networks with homogenous activation functions, either sigmoid (*σ*) or ReLU at every layer, struggled to learn the task. Though, subsets of architectures performed significantly better than the fully recurrent architecture. This supports our use of distinct activation functions per layer across our models.

At half our normal hidden layer size, the average fitness (across all architectures) decreased slightly. But we would draw the same conclusion; in this task, all architectures perform equivalently, or significantly worse than the fully recurrent architecture.

At twice our normal hidden layer size, the average fitness increased slightly. And many architectures perform significantly better than the fully recurrent architecture. This strengthens our case for the benefits of partial recurrence, and suggests that future work, on larger partially recurrent networks, may discover even more benefits.

**Figure 7:**
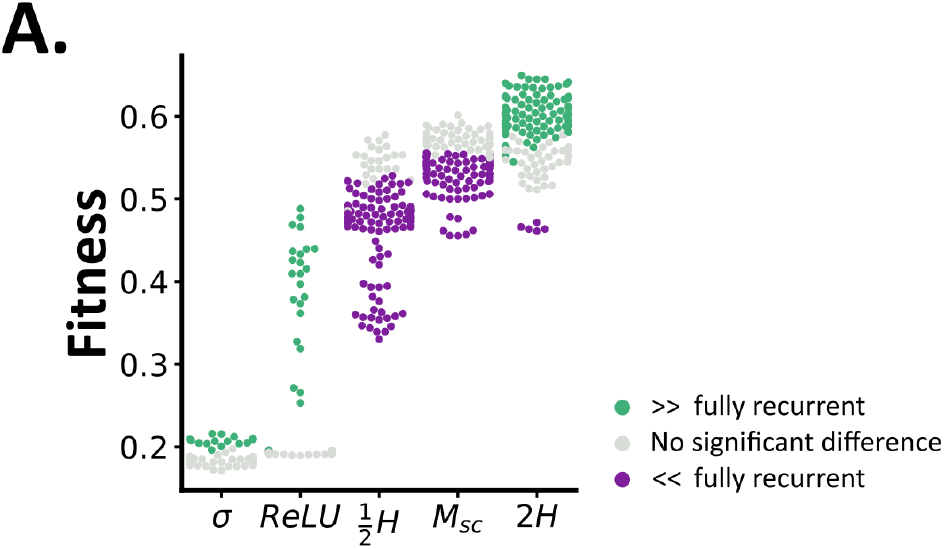
**A** Fitness as a function of hyperparameter configuration. Each circle shows the median fitness, across 50 trained networks, for a single architecture. Architectures which perform significantly better or worse (*p <* 0.01) than the fully recurrent architecture are shown in green and purple, the rest are shown in grey. Results from our typical setup are marked as *M*_*sc*_, *σ* and ReLU indicate networks trained with homogenous activation functions, and ^1^*/*_2_*H* and 2*H* show results from networks with half or twice our normal hidden layer size.

## 7 Acknowledgements

MG is supported by the Eric and Wendy Schmidt AI in Science Postdoctoral Fellowship, a Schmidt Sciences program. We thank members of the Neural Reckoning group, Karim G. Habashy and Jeffery S. Bowers for their constructive input.

